# Genome-wide analysis of FSHD cell lines using Nanopore sequencing reveals allele-specific differences at DUX4 target genes and complex repeats

**DOI:** 10.64898/2025.12.22.696056

**Authors:** Jasmine Shaaban Sakr, Xiangduo Kong, Negar Mojgani, Erisa Taghizadeh, Elnaz Abdollahzadeh, Kyoko Yokomori, Ali Mortazavi

## Abstract

Facioscapulohumeral muscular dystrophy (FSHD) is linked to a monoallelic contraction of primate-specific 3.3kb D4Z4 macrosatellite repeats on the disease-permissive chromosome 4q (4qA haplotype) with additional mutations of a chromatin regulator SMCHD1 acting as a disease modifier. DNA hypomethylation at the D4Z4 repeat and resulting abnormal derepression of the embryonic transcription factor DUX4 encoded in the D4Z4 repeat are the hallmark of FSHD. In order to investigate the impact of FSHD mutations within as well as outside of the disease loci, we performed Nanopore direct-RNA and genomic sequencing to characterize global and D4Z4-specific changes in isoform expression and DNA methylation using CRISPR-engineered human skeletal myoblast lines carrying FSHD mutations (D4Z4 contraction and SMCHD1 mutation) compared to the isogenic parental healthy control line. Nanopore sequencing allowed us to characterize the entire unedited control and contracted D4Z4 arrays as well as distinguish differential methylation patterns at the disease locus on chromosome 4qA from those at a nearly identical nonpathogenic D4Z4 repeat arrays on chromosome 10 and disease non-permissive 4qB allele. We observe hypomethylation both at the DUX4 locus and globally in FSHD mutant cell lines in myoblasts as well as in myotubes. DUX4 target gene expression is correlated with promoter hypomethylation. *De novo* haplotype phasing of genomic and RNA reads reveals allele- and isoform-specific expression of DUX4 target genes as well as highly expressed DUX4 target pseudogenes that may contribute to disease pathogenesis. Taken together, our results indicate significant impact of FSHD mutations not only on D4Z4 allele, but also DUX4 targets and repeat regions in the genome, which may be collectively contributing to the FSHD pathogenesis.

## Introduction

Muscular dystrophy encompasses a group of genetic disorders characterized by progressive muscle weakness and degeneration. Facioscapulohumeral muscular dystrophy (FSHD) is the third most prevalent form of muscular dystrophy, affecting approximately 1 in 20,000 individuals world-wide (1, 2). It is characterized by asymmetric muscle weakness, primarily affecting the face, shoulders, and upper arms, and often leads to disability over time (1). FSHD is subdivided into two types, FSHD1 and FSHD2 (3, 4). FSHD1 (MIM 158900) accounts for approximately 95% of cases and is caused by the contraction of the subtelomeric D4Z4 macrosatellite repeat array (1-10 copies) on chromosome 4q35. In contrast to healthy individuals, the D4Z4 array contains 11 to 100 repeats. FSHD2 (MIM 158901) accounts for 5% of cases with modestly contracted D4Z4 repeats (8-20 copies) and is mostly linked to mutations in SMCHD1 (5, 6), which is an epigenetic modifier (7, 8). Mutated SMCHD1 has also been linked to severe cases of FSHD1, thus acting as a disease modifier gene (9, 10).

D4Z4 is a 3.3 kb macrosatellite repeat containing an open reading frame for the double-homeobox transcription factor *DUX4* gene (11–13). Two subtelomeric variants distal to the D4Z4 repeat array on chromosome 4 have been identified, 4qA and 4qB alleles, and are distinguishable by a pLAM sequence and β-satellite repeat distal to D4Z4 on 4qA (14, 15). Repetitive DNA elements, such as macrosatellites, make up more than half of the human genome (16) and are typically packaged into constitutive heterochromatin. DNA methylation at heterochromatin plays a critical role in controlling gene expression and repressing repetitive DNA elements. Though genetically distinct, both FSHD1 and FSHD2 patient cells exhibit DNA hypomethylation at the D4Z4 region contributing to the expression of full-length double homeobox 4 transcript (*DUX4fl*) (2, 17). Only those individuals with poly(A) signal immediately downstream of the last copy of D4Z4 repeat (4qA permissive haplotype) develop FSHD, suggesting the significance of the specific *DUX4* mRNA production in FSHD pathogenesis (13, 15, 18). Aberrant expression of *DUX4* is linked to activation of gene pathways related to muscle pathology (19–21).

Though DUX4 upregulation is linked to the disease, it is expressed in less than 1% of patient myoblasts and 3-4% of patient myotubes *in vitro* (22–25). How this limited DUX4 expression drives the disease only in patient myocytes and how dysregulation of target genes contributes to the disease process remain obscure and are still under active investigation (17, 23, 26–29). Overexpression (OE) of the recombinant DUX4 in *in vitro* myoblasts and in *in vivo* model organisms is highly toxic (19, 30), which was thought to be a mechanism of the disease. Using single nucleus RNA-sequencing (snRNA-seq) and high-resolution high-throughput spatial transcriptomics (MERFISH), however, we demonstrated that there is no significant sign of cytotoxicity with DUX4 target gene-expressing myotubes in the context of the endogenous DUX4, and that even those seemingly DUX4-negative patient myotubes have undergone transcriptomic changes (31, 32). Our recent results *in vitro* provided evidence that myotubes at late stage differentiation increases DUX4 target expression without toxicity and that the limited DUX4 signal is amplified by a feedforward mechanism of target gene amplification, highlighting the significance of DUX4 target transcription regulators (33).

While many diseases including muscular dystrophies have been studied in animal models such as mice, FSHD is particularly difficult to model because D4Z4 repeats are primate-specific. Furthermore, mice lack two major DUX4 target transcription factors, DUXA and LEUTX (34), and the exogenous *DUX4* expression affects target genes in different signaling pathways in mice and humans (35), indicating the limitation of mouse models to recapitulate the human FSHD phenotype (36). Patient-derived myoblasts serve as an alternative model system, offering insights into DUX4 expression and FSHD-related changes in human cells. However, their limited availability and genetic heterogeneity have been a major obstacle (36). Thus, we recently generated mutant myoblast lines carrying D4Z4 contraction and/or SMCHD1 mutation from healthy human skeletal myoblast line with a permissive 4qA haplotype using CRISPR-Cas9 (33). We demonstrated the synergistic effect of both mutations in heterochromatin disruption and DUX4 target gene activation, highlighting the correlation between heterochromatin loss, DUX4 target activation and disease severity (33).

In the current study, we used the Oxford Nanopore platform genomic DNA and RNA sequencing from our double mutant and parental isogenic control cell lines at the late myotube stage to investigate the epigenomic and gene expression consequences of FSHD mutations in the isogenic background. Our results revealed genome-wide, in some case allele-specific, methylation changes at DUX4 target genes and repetitive elements beyond the alterations observed in the 4q and 10q loci. Furthermore, the robust mappability of long reads allowed detection of allele-specific or isoform-specific DUX4 target gene expression as well as previously underappreciated non-coding homologs that may contribute significantly to the FSHD phenotype. Taken together, our Nanopore sequencing approach revealed intricate genome-wide epigenetic and gene expression changes induced by FSHD mutations in the context of the endogenous *DUX4* at higher resolution than previously appreciated with short-read sequencing.

## Results

### Detection of genome-wide methylation in control and isogenic FSHD mutant cells

The CRISPR mutant human skeletal myoblast line contains a single repeat unit on the 4qA allele and 2 repeat units on the 4qB allele with SMCHD1 null mutations on both alleles (Fig. 1A) (33). The isogenic parental control and mutant myoblast lines (termed “FSHD mutant” or “FSHD” hereon) were cultured and differentiated into late myotube stage on days 12 and 14, respectively. Compared to the early myotube stage (differentiation days 3-4), differentiation marker genes such as *MYH1* and *MYL3* are robustly upregulated at a significantly higher level at the late myotube stage (33). Duplicates of genomic samples (total of four) corresponding to undifferentiated proliferating myoblast stage and differentiated late myotube stage of control and FSHD myocytes were sequenced using Oxford Nanopore for an average of 60 gigabases (Gbs) per sample and a minimum coverage of approximately 20x (Fig. 1B). RNA from control and FSHD myotubes were also sequenced using direct-RNA sequencing in biological duplicates to an average of 10 millions reads per replicate. Reads were mapped using the Dogme pipeline (37) using both the GRCh38 and the CHM13 T2T assemblies to extract DNA methylation (Methods). Most analyses in the current study unless otherwise noted are based on the CHM13 T2T assembly. Each sample had at least 50 million sites with at least one modification and 43.3 million (around 83%) are the same sites in all four samples. Of these genome-wide sites, an average of 98.86% are methylated (5mC) and 21.18% are hydroxymethylated (5hmC) (Fig. 1C). The low detection of hydroxymethylation is consistent with the fact that muscle is not a tissue enriched in the 5hmC modification (38, 39).

**Fig. 1.**
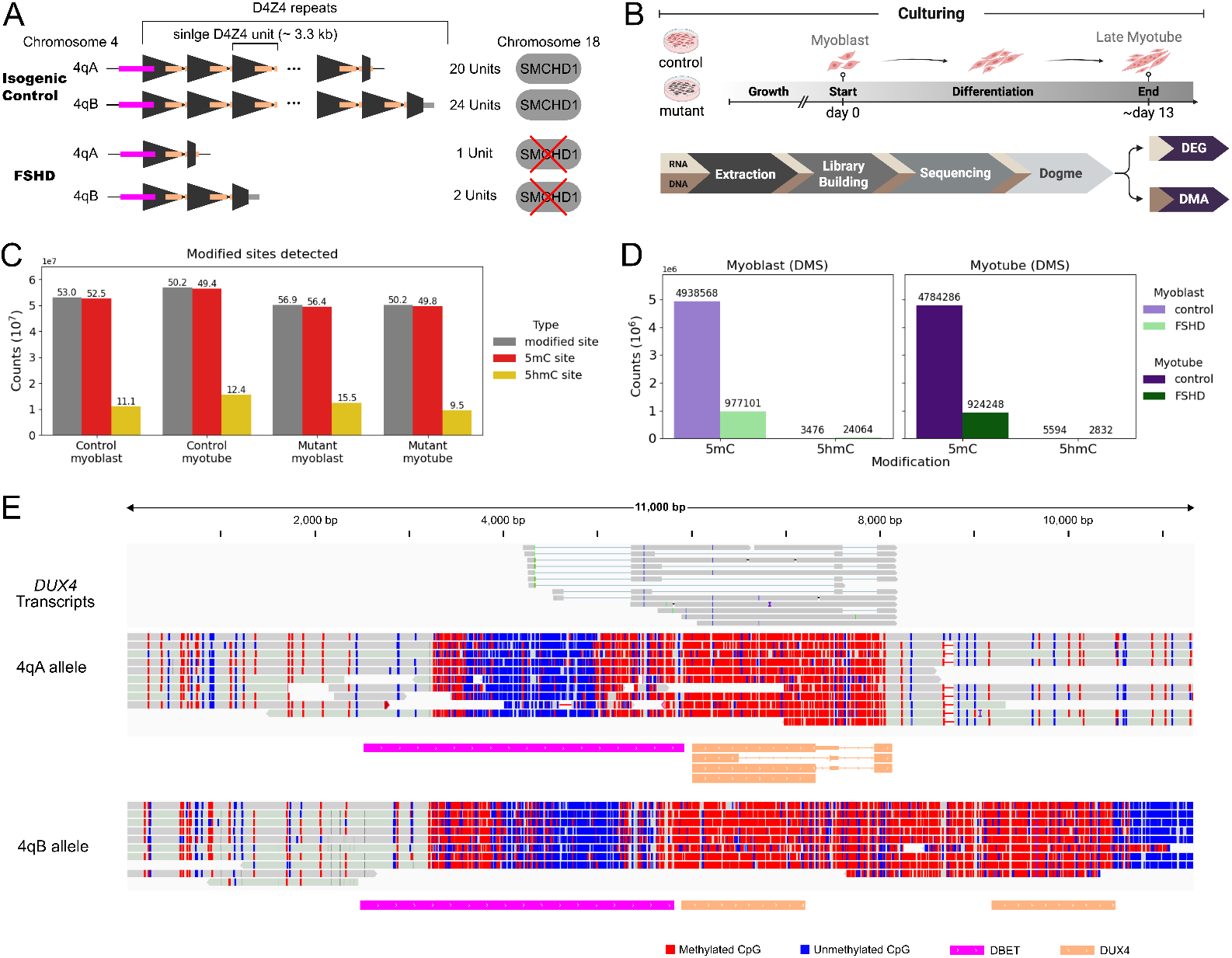
Hypomethylation detected genome-wide and at disease locus in FSHD mutant cell line compared to isogenic control. **A**. D4Z4 repeat status and *SMCHD1* mutation in control and CRSPR-edited FSHD mutant myoblast lines. In the control, the permissive 4qA allele has 20 repeat units and the 4qB allele has 24 repeats units. The FSHD mutant cell line has a single repeat unit on the 4qA allele and 2 repeat units on the 4qB allele. In the mutant cells, both copies of SMCHD1 are mutated. **B**. Overview of experimental and analysis workflow. **C**. Each sample has over 50 million sites with modifications (grey) after filtering out sites at low coverage. The majority of these sites have 5mC modification (red) and only a fraction have 5hmC modification (yellow). **D**. Mutant cells are hypomethylated at differentially methylated sites (DMS). In both myoblast and myotubes, over a million 5mC DMS (p-value 0.05) were detected and a majority were hypermethylated in control (light and dark purple), but not in the FSHD mutant cells (light and dark green). **E**. FSHD late myotube stage genomic reads span the entire locus and allow for accurate characterization of repeat units. Hypomethylation at the *DUX4* promoter is observed as well as full-length *DUX4* transcripts with different isoforms detected, including unspliced transcripts.

### Global DNA methylation changes in FSHD mutant cells at myoblast and myotube stages

Next we compared the methylation level per site between FSHD mutant and control in both undifferentiated myoblasts and differentiated late myotubes. There is an overall significant difference in methylation level between FSHD mutant and control at these same shared 43.3 million sites in both myoblasts (Mann-Whitney statistic 1.01×10^15^, p-value < 1 x 10^-308^) and at the late myotube stage (Mann-Whitney statistic 9.32×10^14^, p-value < 1 x 10^-308^). In myoblasts, 5,915,669 sites are 5mC differentially methylated sites (DMSs) (Methods) and 83.48% are less methylated in FSHD mutant than control (Fig. 1D). In the late myotube stage, 5,708,534 sites are 5mC DMSs and 83.81% are less methylated in FSHD mutant than control. Around 37% of the 5mC DMSs are the same site at both stages and over 76% of 5mC DMSs in myoblasts are at or within 2 base pairs of a corresponding 5mC DMS in late-stage myotubes. These findings indicate a significant global hypomethylation in FSHD mutant compared to isogenic control, which is driven by a reduction in methylation level rather than loss of methylated sites. This difference is maintained during differentiation into the late myotube stage, as the majority of the DMSs are in the same loci and exhibit hypomethylation. This is further reflected at CpG Islands genome-wide. There is also a significant difference in methylation level in myoblasts (Mann-Whitney statistic 1.41×10^9^, p-value 4.23 x 10^-98^) and in late myotube stage (Mann-Whitney statistic 1.35 x 10^9^, p-value 6.07 x 10^-22^). Mean level methylation is lower in FSHD mutants (T statistic 5.13, p-value 2.87 x 10^-7^) than control (T statistic 3.23, p-value 1.22 x 10^-3^), indicating hypomethylation at CpG islands. Overall, there is a global 5mC hypomethylation in FSHD mutant cells and remains relatively hypomethylated across muscle differentiation.

### Allele-specific expression of *DUX4fl* transcripts correlating with differential DNA hypomethylation and haplotypes

We observe hypomethylation at the D4Z4 locus in the FSHD mutant cells (Fig. 1E) as described previously (Kong et al., 2024). The proximity of *DBET* to *DUX4* in FSHD mutant myotubes as a result of repeat contraction creates a haplotype-specific promoter on 4qA. The *DUX4* promoter region, defined as 2kb upstream and 500 base pairs (bp) downstream of the transcription start site (TSS), on the permissive 4qA allele (3) is 47.22% methylated compared to 81.31% methylated on the non-permissive 4qB allele, which are respectively 75.98% and 63.87% in controls (Fig. 1E, Fig. S1A. We specifically detected monoallelic expression of *DUX4* transcripts from 4qA, but not from non-permissive 4qB allele. Several of the 14 reads containing the full-length 4qA *DUX4* ORF detected showed unique splicing patterns retaining introns (Fig. 1E). We also measured methylation at the highly-homologous, nonpathogenic D4Z4 repeats on chromosome 10q (where alleles are denoted as either H1 or H2). The control is methylated on both alleles (H1 79.5% methylated and H2 74.70% methylated) (Fig. S1B). The CRISPR editing resulted in repeat contraction on chromosome 10q in the FSHD mutant cell line (Kong et al., 2024) (33). Interestingly, while one allele is hypomethylated, the other allele kept intact methylation as in the control (H1 46.57% methylated and H2 78.99% methylated) (Fig. S1B). Despite the genomic and DNA methylation changes, no RNA reads were mapped to the *DUX4* gene on chromosome 10. Thus, our results indicate specific *DUX4* full-length (*DUX4fl*) transcript expression, including some splicing variants, from 4qA, but not from 4qB or 10q.

### Upregulation of DUX4 target genes is accompanied by promoter hypomethylation

We detected 16,396 genes expressed in one or more of our mutant and control samples. We examined 63 DUX4 target genes that are upregulated in FSHD patient and mutant cells and were shown to have *DUX4* binding to their promoter regions by ChIP-Seq (32– 34). As expected, we detected a significant upregulation of these DUX4 target genes at the late myotube stage (Fig. 2A) with 25 of the 63 significantly upregulated (p-value 0.05). The top 10 differentially expressed DUX4 target genes were *ZSCAN4, CCNA1, TAF11L11, TRIM43, H3Y1, TAF11L12, SLC34A2, TRIM43B, KHDC1L*, and TRIM49C (Table S1). We examined whether activation of these genes is accompanied by any change in DNA methylation at the promoter regions (Fig. 2B). Since only 13 of the DUX4 target genes have annotated promoters in ENCODE cis-regulatory elements (cCREs) (40), the methylation status was measured at a 2.5 kb promoter spanning region for the DUX4 target genes as well as other FSHD associated genes *DUX4c, FRG1* and *FRG2*.

**Fig. 2.**
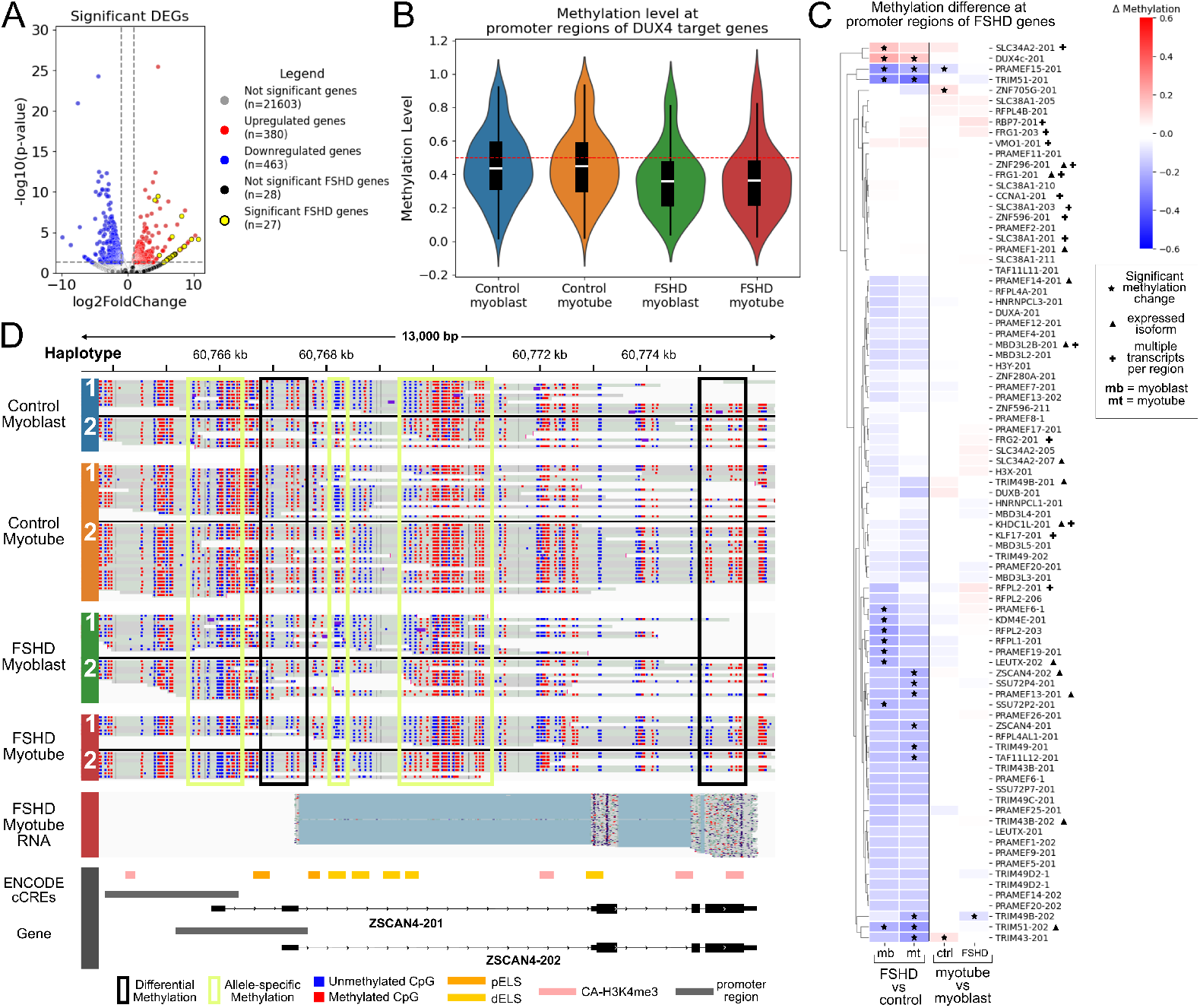
Significant hypomethylation and upregulation of DUX4 target genes in FSHD mutant cells. **A**. Of 63 DUX4 target genes, 25 were significantly upregulated (p-value 0.5) (yellow). **B**. Mean methylation level at 2.5kb promoter regions of DUX4 target genes. The mean is lower in FSHD mutants in both myoblasts (green) and myotubes (red). **C**. Methylation difference at DUX4 target gene promoter regions between (left to right) FSHD mutant and control myoblasts, FSHD mutant and control myotubes, control myotubes and myoblasts, and FSHD mutant myotubes and myoblasts. Significant methylation, isoform expression and promoter regions with multiple isoforms are annotated with a star, triangle, or a plus (respectively). **D**. Genomic reads and FSHD mutant myotube RNA reads at the *ZSCAN4* locus. Allele-specific (yellow box) and differential methylation (black box) are indicated.

*DUX4c* has been shown to have an antagonistic effect on *DUX4* and downregulated its targets (41). FSHD region gene 1 *(FRG1)*and FSHD region gene 2 (*FRG2)* are both upregulated in FSHD and has been implicated in muscle differentiation (42–45). Overall, FSHD has a mean methylation of 37.2% compared to 45.6% in control at DUX4 target genes (Fig. 2B). On average, 78% of DUX4 target genes promoter regions are hypomethylated in FSHD, 19% exhibit no change (Methods) and 3% are hypermethylated in FSHD (Fig. 2C). *DUX4c* is significantly hypermethylated (16.64% methylated in control, 37.78% methylated in FSHD) and no corresponding transcripts were detected in the late myotube stage. There is no significant DNA methylation at the promoter regions of *FRG1* and *FRG2* in both control and FSHD mutant myotubes, and while FRG1 is expressed in both, no significant *FRG2* transcripts were detected in either control or FSHD mutant cells at this stage (Table S1).

We found that the DUX4 target genes exhibiting the strongest differential expression show reduction in methylation in the promoter regions of approximately 15% or more in the FSHD mutants. In the top 10 differentially expressed, these include *ZSCAN4* (68.06% methylated in control, 48.29% methylated in FSHD, average of isoforms), *TAF11L12* (58.46% methylated in control and 40.17% methylated in FSHD) and *TRIM43* (80.93% methylated in control, 56.79% methylated in FSHD). We also observed another subset of DUX4 targets and are similarly significantly hypomethylated but resulted in low expression in FSHD mutant cells. The most striking example is *TRIM51* (47.10% methylated in control, 17.17% methylated in FSHD, average of two isoforms) where the reduction in methylation is around 30% but is expressed at less than 2 transcripts per million (TPM). In contrast, the *TRIM43* gene exhibits the reduction in methylation at the promoter is around 24% and is expressed at over an average of 20 TPM. Other genes with more than 15% reduction and less than 2 TPM expression include *PRAMEF13, PRAMEF15, RFPL1, RFPL2*. This indicates that although there appears to be a significant correlation between activation of DUX4 target genes and promoter hypomethylation, not all the genes with high reduction in methylation were significantly expressed at this late myotube stage in muscle differentiation. Thus the correlation may in part be dependent on the stage of differentiation.

### Isoform-level analysis reveals a subset of preferentially expressed and differentially methylated DUX4 target genes

Long-read sequencing enables detailed investigation of isoform expression. However, characterizing DUX4 target isoform expression remains limited. We focused on analyzing known transcripts as well as novel transcripts that were detected in at least two samples with a minimum of 0.1 TPM expression in two or more samples using reconcileBams (Methods). We detected 45,142 known transcripts and 127,134 novel transcripts. We detected 24 DUX4 target genes that have more than one isoform (protein-coding and processed transcript) and 17 of these possess isoforms with distinct TSS, resulting in non-shared promoter regions. In late myotube stage, we found that 13 of DUX4 target genes exclusively express only one isoform, 4 of which are included in the top 10 differentially expressed DUX4 target genes in late myotube stage, including *ZSCAN4, SLC34A2, TRIM43B* and *KHDC1L* (Fig. 2C, D). Of the 13 DUX4 target genes, 10 of them contain 2 or more distinct promoter regions, most of which are hypomethylated in FSHD mutant cells, with isoforms from the same gene exhibiting comparable promoter methylation levels (Fig. 2C, Fig. 2A). A few genes have significant differences in methylation in the promoter region of an isoform that is not expressed. Although the promoter of *TRIM49B-202* is significantly hypomethylated (49.59% methylated in control, 30.21% methylated in FSHD), the *TRIM49B-201* isoform is expressed. Similarly, while the promoter of *PRAMEF1-202* is significantly hypomethylated (35.51% methylated in control, 22.99% methylated in FSHD), the *PRAMEF1-201* isoform is expressed. *SLC34A2* is one of the few genes with a significantly hypermethylated promoter (37.08% methylated in control and 53.39% in FSHD). This promoter appears to be utilized by 5 of 7 isoforms, but the significantly expressed transcript, *SLC34A2-207*, does not use this promoter. These observations reveal examples of discordance between DNA methylation change in the promoter region and corresponding isoform expression, and suggest potential local regulatory elements acting specifically on the expressed isoform. They also highlight the need for higher-resolution methylation profiling at isoform-specific alternative promoters rather than across a broad promoter region. Nevertheless, we have begun to characterize isoform-specific expression patterns of DUX4 target genes.

### DNA hypomethylation at DUX4 target gene promoters is observed in myoblasts prior to robust activation in myotubes

We observed a similar trend of hypomethylation at DUX4 target promoters in FSHD mutant cells at both myoblast and late myotube stages even though many of the DUX4 target genes are not highly induced until after myotube differentiation (Fig. 2C) (33). A subset of *PRAMEF* genes and *TRIM51* were significantly hypomethylated before muscle differentiation. Other DUX4 target genes were more hypomethylated in myoblasts than in the late myotubes stage. These include *RFPL2*-203 (−24.2740 mean difference in myoblast and −17.32 mean difference in myotubes), *RFPL1* (-21.99% mean difference in myoblast and 17.25% mean difference in myotubes and *KDM4E* (−19.80 mean difference in myoblast and −10.40 mean difference in myotubes), all of which are lowly expressed in late myotube stage. The promoter region of the *LEUTX*-202 isoform is also more hypomethylated in myoblasts (−20.03 mean difference in myoblast and −15.07 mean difference in myotubes) but is significantly expressed in the late myotube stage. Thus, our results indicate that overall, the promoter regions of the DUX4 target genes are already hypomethylated in the proliferating myoblast stage, and mostly remain hypomethylated into the late myotube stage. This raises the possibility that a low level *DUX4* signaling in undifferentiated myoblasts is sufficient to open up the target gene promoters, contributing to the effective upregulation of these genes in differentiated myotubes. It should be noted, however, that not all methylation loss leads to expression in myotubes, indicating that DNA methylation is not the sole determinant of gene expression changes in FSHD mutant cells.

### Subset of DUX4 target pseudogenes are highly expressed

Long-read RNA-seq also enables the ability to distinguish reads originating from a gene versus its pseudogene(s). In particular, we were able to map and define the significant upregulation of two classes of pseudogenes in FSHD mutant cells at more than 2 (TPM) with no expression in control in myotubes. Many of these were not detected or not properly mapped using short-read sequencing (33). The pseudogene with the highest expression is the histone H3 variants/pseudogenes in the locus of the TAF11-like microsatellite repeat array on chromosome 5. Previously, 3 H3-like genes in this locus, histone variants *H3*.*X (H3Y2) and H3*.*Y (H3Y1)* as well as a pseudogene *H3*.*Z (H3P21)*, were reported to be upregulated in DUX4-induced myoblasts (46). In our FSHD mutant late myotubes, *H3*.*Y (H3Y1)* is also among the top 10 differentially expressed genes while *H3*.*X (H3Y2)* is very lowly expressed (0.6 average TPM). The DUX4 target pseudogene with the highest expression in FSHD mutant cells is H3.Z (*H3P21)* (p-value 2.4 x 10^-5^, 82.39 average TPM) (Table S1). Indeed, *H3*.*Z* has a 13.33% reduction in methylation at the promoter, compared to only 3.59% reduction at the *H3*.*X* promoter, in FSHD mutant myotubes (Fig. 3A). Another less characterized H3 histone pseudogene, *H3P18*, in this locus was also significantly expressed (p-value 1.38 x 10^-3^, 16.32 TPM). Hypomethylation at the *H3P18* promoter is more pronounced in the myoblasts (- 12.08% mean difference) than in the late myotube stage (- 3.94% mean difference). *H3P18* has 83.95% sequence identity with the aligned region at *H3*.*Y* (42% alignment coverage), similar to *H3*.*Z* sequence identity (84.49%) with the aligned region at *H3*.*Y* (37% alignment coverage). Neither *H3*.*Z* nor *H3P18* was detected in short-read data. Thus, our results indicate that expression of multiple H3-related variants/pseudogenes in this locus is affected by FSHD mutations.

**Fig. 3.**
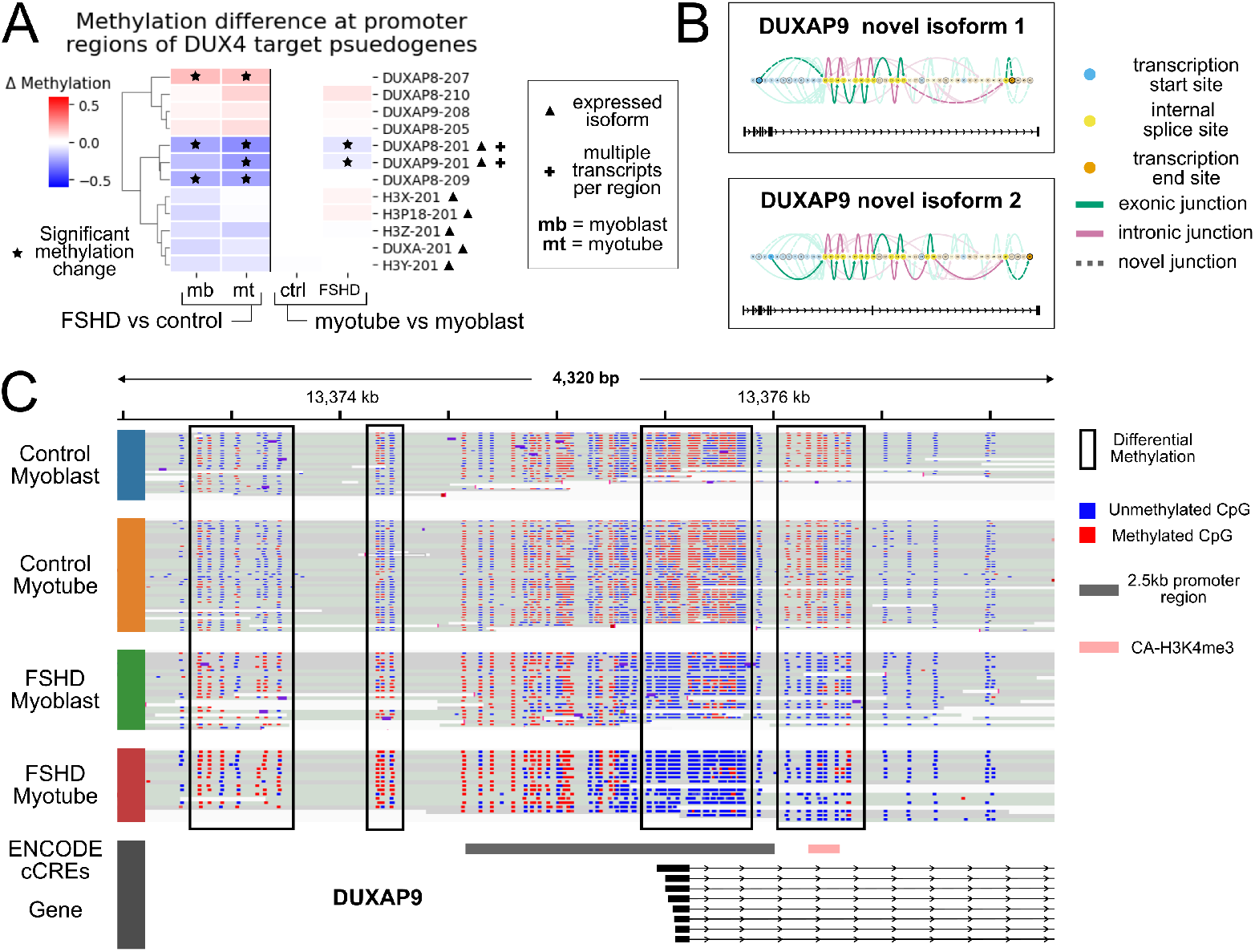
Significant hypomethylation and upregulation of DUX4 target pseudogenes in FSHD mutant cells. **A**. Methylation differences at DUX4 target pseudogene promoter regions between (left to right) FSHD mutant and control myoblasts, FSHD mutant and control myotubes, control myotubes and myoblasts, and FSHD mutant myotubes and myoblasts. Significant methylation, isoform expression and promoter regions with multiple isoforms are annotated with a star, triangle, or a plus (respectively). **B**. Swan graph visualization of novel *DUXAP9* pseudogenes using the swanViewer package. Both are significantly expressed Novel-In-Catalog (NIC) isoforms. **C**. Genomic reads and FSHD mutant myotube RNA at *DUXAP9*. Differential methylation (black box) observed.

*DUXA* is a long-known DUX4 target gene, and was recently shown to suppress other DUX4 target genes (47). We found that two *DUXA* pseudogenes are expressed significantly higher than the actual *DUXA* in FSHD mutant cells at the late myotube stage. Those are *DUXAP9* on chromosome 14 (p-value 7.09 x 10^-4^, 21.32 TPM) and *DUXAP8* on chromosome 22 (p-value 1.58 x 10^-3^, 15.11 TPM) while *DUXA* on chromosome 19 itself was very lowly expressed (around 0.5 TPM) (Table S1). *DUXAP8* and *DUXAP9* both encode long-noncoding RNAs known to play a role as oncogenes in promoting cancer malignancy (48–51), but characterization in the context of muscle disease is very limited (52, 53). *DUXAP9* and *DUXAP8* have 9 and 10 isoforms, respectively, and multiple isoforms have been detected for each, including two significantly expressed Novel-In-Catalog (NIC) isoforms for *DUXAP9* (Fig. 3B). Both of these novel isoforms are expressed at a higher level (both at an average of 3 TPM) than the expressed *DUXAP9-207* (average 2.35 TPM). These *DUXAP9* and *DUXAP8* isoforms each share the same respective promoters, both of which are significantly hypomethylated in FSHD mutant cells, with reductions exceeding those observed at the *DUXA* promoter region (Fig. 3A). In myoblasts, there is a 16.95% reduction in methylation at the *DUXAP9* promoter region (Fig. 3C) and 22.63% reduction at *DUXAP8*. Hypomethylation is increased in the late myotube stage, with a 25.51% reduction at the *DUXAP9* promoter region (Fig. 3C) and 28.53% reduction at *DUXAP8*. This differs from the more minimal change in methylation at *DUXA* promoter (−12.00% mean difference in myoblasts and −8.12% mean difference in late myotubes stage) (Fig. 3A). Given the pronounced hypomethylation at *DUXA* pseudogenes, we revisited our short-read data to assess their expression. Only *DUXAP8* was detected, with an average TPM of 0.19 in controls and 63.21 in FSHD mutant cells. The *DUXA* gene has a 614 bp coding sequence (CDS) and there is a 88% sequence identity with both *DUXAP9* and *DUXAP8*. In addition to the high sequence homology in the CDS, both pseudogenes are located in genomic regions with low unique mappability. Only 24.3% of *DUXAP8* exons and 14.7% of *DUXAP9* exons overlap uniquely mappable 50 k-mers. This is in contrast to *DUXA* which is located completely within a uniquely mappable region. As a result, accurately mapping reads to its pseudogenes using short-read sequencing is likely more challenging. Taken together, our results reveal that long-read sequencing enables more accurate assignment pseudogene transcripts, which was not possible with conventional short-read sequencing, and highlight the significant upregulation of pseudogenes homologous to the protein coding DUX4 target genes in FSHD mutant cells. Expression of pseudogenes, in some cases better than the original gene, by *DUX4* raises the possibility that these pseudogene products (e.g., lncRNAs) may have critical roles in FSHD pathogenesis. It is important to note that we found evidence for reference assemblies also affecting the mapping of these pseudogenes and other DUX4 target genes even with long read data. For a subset of genes, the TPMs show significant discrepancies between GRCh38 (Hg38) (Table S2) and the complete telomere-to-telomere (T2T) human genome assembly (T2T-CHM13v2.0) (Table S1). *DUXAP9* TPM is higher when mapped against Hg38 (average 30.5 TPM) compared to Chm13 (average 21.3 TPM). Another DUXA pseudogene, *DUXAP10*, has a TPM more than doubled in Hg38 (average 13.32 TPM) compared to Chm13 (average 3.48 TPM). The DUX4 target gene *TRIM43B* is lowly expressed when mapped to Hg38. Therefore, mapping RNA-seq reads to Hg38 versus Chm13 can lead to differences in TPM estimates, which affects the interpretation of gene expression and isoform usage.

### Phasing of genomic and RNA long-reads reveal unequal allelic DNA methylation and RNA expression

Long-read sequencing enables us to distinguish two alleles; as long as at least one SNP is anywhere in the read, two haplotypes can be distinguished. We performed *de novo* haplotype phasing to investigate allele-specific expression of DUX4 target genes (Methods). Although approximately half of these genes contained an exonic SNP, most were either not allele-specific or were too lowly expressed to assess reliably. However, phasing of RNA reads identified three isoforms that are either disproportionately expressed from one allele or exclusively expressed from a single allele. *RFPL4A* transcripts are expressed from both alleles (which will be denoted as H1 and H2 with distinct, mostly non-overlapping gene promoter regions counted separately) but with significantly different ratios; 84.21% transcripts are coming from H2 allele (Fig. 4A), correlating with more severe hypomethylation (H1 −3.65% average mean difference and H2 and −11.74% average mean difference). *TRIM49* and *ZNF280A* exhibit monoallelic expression, however *ZNF280A* is in both control and mutant cells. *TRIM49-201* expression originates from the H1 allele that is more broadly methylated across the full 2.5kb promoter region (H1 −24.78% average mean difference and H2 −21.1291% average mean difference) however, closer inspection reveals distinct demethylation patterns on the expressed allele that are not present on the non-expressed allele. *ZNF280A* expression originates from the more hypomethylated H2 allele (H1 0.47% average mean difference and H2 and −11.26% average mean difference) and the 344 bp annotated promoter is nearly completely demethylated on that allele.

**Fig. 4.**
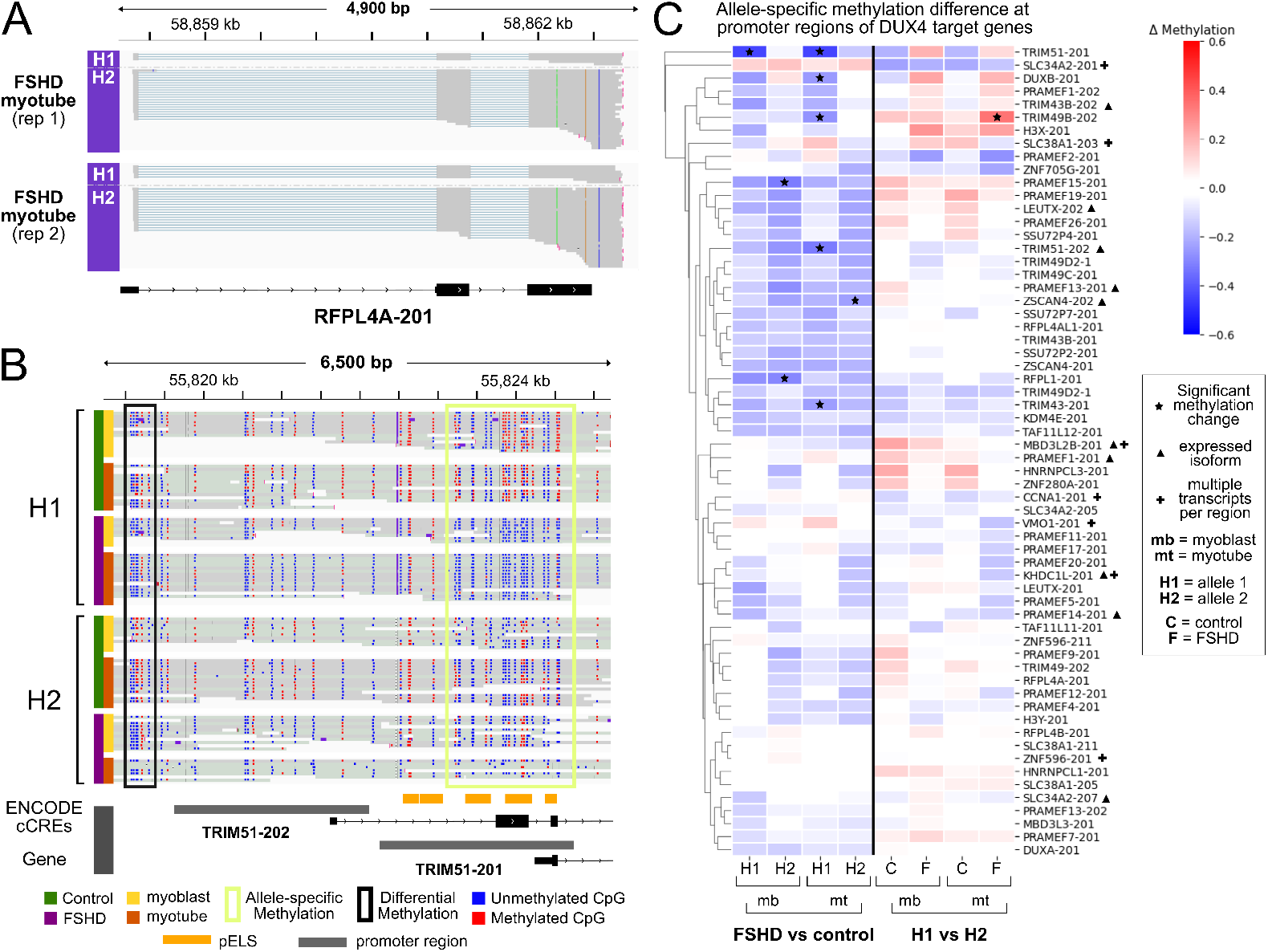
Phasing reveals allele-specific methylation and expression of DUX4 targets. **A**. Phasing using exonic SNPs at *RPL4A* show that the majority of transcripts originate from the H2 allele compared to H1 which has no SNPs. **B**. Genomic reads at *TRIM51*. Top four genomic reads tracks are reads that originated from one allele (H1) and the last four genomic reads tracks are reads that originated from the other allele (H2). Allele-specific (yellow box) and differential methylation (black box) are indicated. **C**. Allele-specific methylation difference (H1 vs H2) at DUX4 target gene promoter regions between (left to right) FSHD mutant H1 and control H1 as well as FSHD H2 and control H2 at myoblast stage (mb) and myotube stage (mt). H2 and H1 comparison in control (C) and FSHD (F) myoblast (mb) and myotubes (mt) stages.

### Phasing reveals allele-specific methylation at DUX4 promoter regions

We also performed *de novo* haplotype phasing to genomic reads to assess whether methylation levels exhibit allele-specific differences at DUX4 target gene promoters. Allele-level analysis revealed that although the majority of genes are hypomethylated in FSHD, methylation levels at promoter regions can differ substantially between alleles (Fig. 4B, C, Fig. S2B). In myoblasts, 77.05% of promoter regions are differentially methylated between control and FSHD mutant on one allele but 90.16% are differentially methylated on the other allele. In the late myotube stage, 78.68% of promoter regions are differentially methylated on one allele and 85.24% are differentially methylated on the other allele. This suggests that some significant differences in methylation between FSHD mutant cells and control could be driven by changes on a single allele rather than the combined changes across both alleles. The most striking example of allele-specific methylation differences in the DUX4 target genes is with *TRIM51*, which exhibits the highest reduction in methylation at its promoter region among the DUX4 targets at around 30% (Fig. 4B, C). We found that the *TRIM51-201* promoter region is significantly hypomethylated only on one allele (*TRIM51-201* H1 −44.33% average mean difference and H2 −09.82% average mean difference). Upon further investigation, we observe a near complete demethylation at a 348 basepair candidate proximal enhancer at the *TRIM51-201* TSS in the significantly hypomethylated allele in FSHD (Fig. 4B). Despite these allelic methylation patterns associated with *TRIM51-201*, the second isoform, *TRIM51-202*, is the one that is actually expressed in FSHD mutant myotubes. *DUXB* is another DUX4 target for which phasing reveals a distinct allelic pattern that is obscured in the collapsed analysis. The hypomethylation at the promoter region in myoblast is not significant (−5.96% mean methylation difference) compared to the late myotube stage (−15.03% mean methylation difference). However, allelic methylation analysis reveals that the one allele is significantly hypomethylated (H1 −25.23% average mean difference compared to H2 −5.64% average mean difference) at both stages of differentiation. Similarly to *TRIM51-201*, there is also a near complete demethylation at a 335 base pair candidate annotated promoter of *DUXB* in the significantly hypomethylated allele. Overall, most alleles are hypomethylated in FSHD mutant cells but at varying levels. The promoter regions with hypomethylation on both alleles (regardless of magnitude) drop to 54% in myoblasts and 60% in the late myotube stage from 78% in the bulk analysis. Our results reveal complex DNA methylation patterns within each individual allele, which may play an important role in DUX4 target gene expression. The results further highlight the need to perform high-resolution DNA methylation analyses down to specific regulatory elements in individual alleles.

### FSHD mutations allele may contribute to allelic-methylation differences at promoter regions

We compared methylation levels between the two H1 and H2 alleles at DUX4 target gene promoters within each sample and then compared these patterns (H1 vs H2) between the control and FSHD lines to determine whether FSHD mutations altered allele-specific methylation (Fig. 4C). In myoblast, the proportion of DUX4 target promoters with allele-specific methylation are 78.68% of promoters in control and 83.60% of promoters in FSHD, and the two alleles do not necessarily change in the same direction (e.g., both becoming hypomethylated). In myotubes, although the proportion of DUX4 target gene promoter regions with allele-specific methylation are comparable between control and FSHD mutant cells (72.13% of promoters in both), the directions of the changes in mutant cells are different from those at the myoblast stage. For example, 34.42% of promoters are more hypomethylated on one allele in control myotubes compared to myoblast stage while 47.54% of promoters are hypomethylated on one allele in FSHD mutant cells. It is important to note, that *de novo* phasing does not allow to definitively determine whether the more hypomethylated promoters are inherited from the same parent as the permissive allele. However these observations suggest that although there are differences in methylation between the two alleles in the control, FSHD mutations may have further exacerbated these differences between H1 and H2 alleles.

### Hypomethylation is also associated with *DUX4*-mediated transcription of retroelements and satellite repeats

In addition to heterochromatin loss at D4Z4 repeat regions, *DUX4* has been shown to reactivate transposable elements, which may also be associated with epigenetic changes (54–56). Thus, in addition to unique gene regions, we examined methylation levels of all repeat classes in the genome. In both myoblast and myotubes, FSHD mutant cell line has a lower mean methylation (59.06% and 60.99%, respectively) than control (65.35% and 66.31%, respectively) across all the repeats in the genome (Fig. 5A). Satellite repeats have the lowest mean methylation in the mutant cells (p-value 5.2 x 10^-3^), in both myoblasts (−10.87% mean difference) and myotubes (−11.05% mean difference), compared to controls followed by long terminal repeat (LTR) elements (p-value 3.5 x 10^-2^, −6.77% methylation difference in myoblast and −6.05% methylation difference in myotubes) (Fig. 5B). This is consistent with the activation of retroelements as well as satellite repeats by *DUX4* in both FSHD patient myocytes and during the embryonic genome activation (EGA) (55, 57). In agreement with *DUX4*-induced expression of transcripts derived from ACRO1 and HSATII satellite repeats (58), ACRO1 and HSATII had the lowest mean methylation in FSHD mutant cells (−13.91% and −12.66% respectively, average between myoblast and myotube) (Fig. 5C). Taken together, DNA hypomethylation not only associates with unique DUX4 target gene promoters, but also with repetitive elements.

**Fig. 5.**
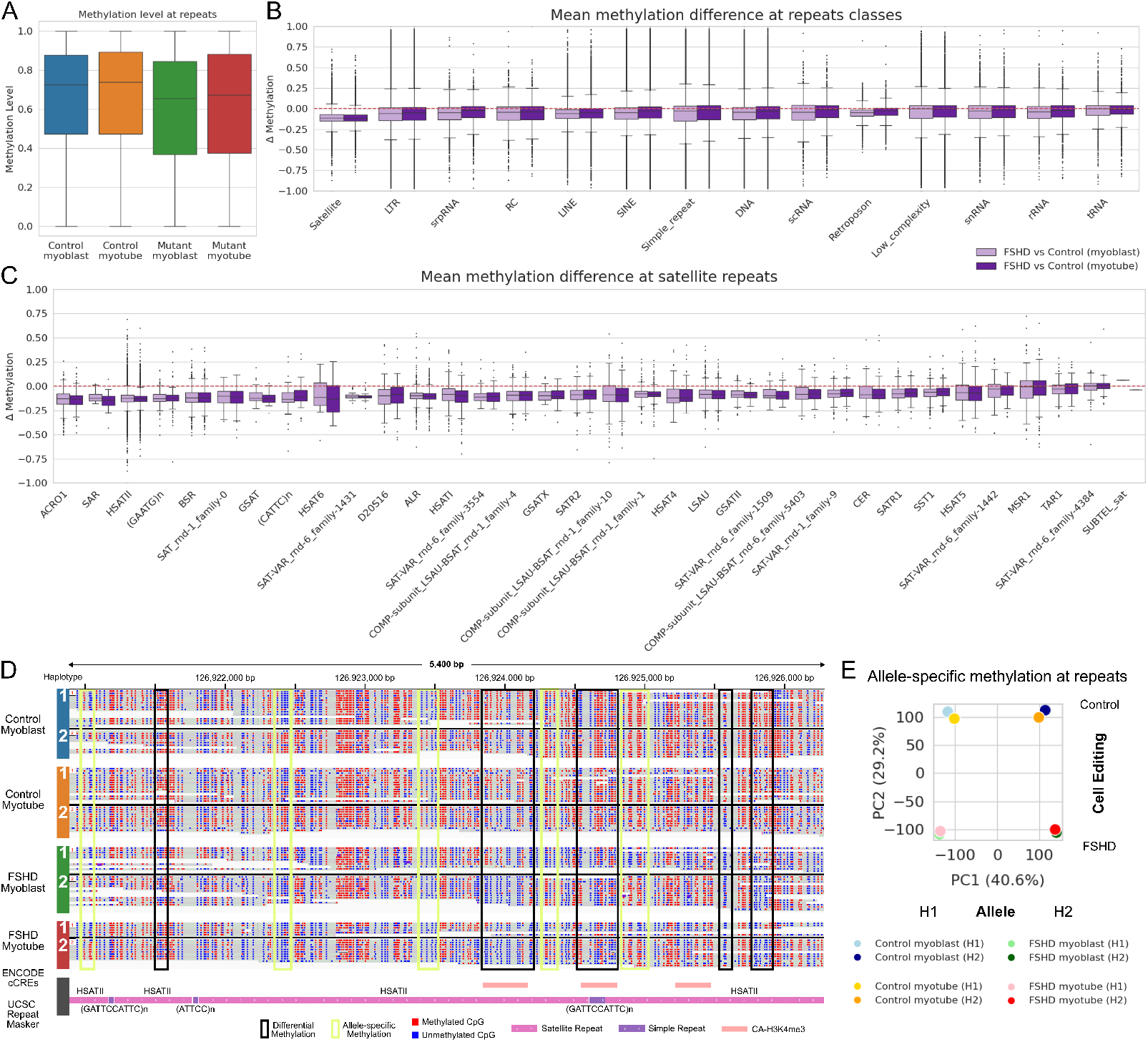
Hypomethylation in satellite repeats and retroelements. **A**. Mean methylation level of repeats genome-wide is lower in FSHD mutants in both myoblasts (green) and myotubes (red). **B**. Mean methylation of alpha, beta and other satellites including D4Z4-like repeats across the genome are lower than other repeat classes, followed by LTR. **C**. The satellite repeats with the lowest mean methylation are acrocentric, SAR and HSATII. D4Z4 repeats are classified under the COMP-subunit_LSAU-BSAT. **D**. Methylation of each cell line shows sample-specific (black boxes) and allele-specific (yellow boxes) methylation both at the promoter region and across the repeats. **E**. There is more variance in methylation levels between alleles at repeats (PC1 40.6%) than between control and mutant (PC2 29.2%).

Based on the allele-specific methylation patterns observed at DUX4 target promoters, we next asked whether similar patterns were present at repeat elements. After phasing, we did observe allele-specific methylation at repeats (Fig. 5D). PCA reveals that allelic identity accounts for the largest source of variation (Fig. 5E). PC1 (40.6%) separates the two alleles, where positive PC1 is H2 and negative PC1 is H1. Despite being phased without parental genotype information, the random haplotype assignments revealed substantial methylation differences, which emerged as the largest source of variance across all repeat elements. The second largest source of variance (PC2 29.2%) is driven by FSHD mutations, positive PC2 is control and negative PC2 is FSHD. These observations further support the potential impact of the pathogenic allele on methylation patterns genome-wide.

## Discussion

We used long-read genomic DNA and RNA sequencing to characterize genome-wide epigenetic effects of FSHD mutations. Long-read sequencing enabled resolution of complex genomic regions, transcript detection of isoforms/homologs and allelic imbalance. The Oxford Nanopore platform extends this capability by allowing concurrent identification of structural variants and nucleotide modifications on individual reads (59, 60). Nanopore sequencing is increasingly being adopted in FSHD research because it allows simultaneous molecular classification of samples (61, 62) and direct measurement of methylation changes (60, 63). This reduces the need for additional material for utilizing multiple separate assays such as optical mapping (64, 65) and short-read, bisulfite sequencing (66, 67) for the same genomic and epigenomic information, which is advantageous given the limited availability of patient muscle biopsies. However, the majority of the studies using Nanopore focus on the 4q35 and 10q loci (68–70) or D4Z4 and D4Z4-like repeats genome-wide (71). We leverage Nanopore sequencing to characterize effects FSHD mutations in double mutant (i.e., D4Z4 repeat contraction and SMCHD1 mutation) and parental isogenic control cell lines, enabling more confident assessment of epigenetic and transcriptomic changes driven by FSHD independent of background genetic variation.

In this study, we found significant global 5mC hypomethylation in the FSHD mutant cells compared to isogenic control where the hypomethylation reflects lower methylation levels rather than a decrease in the number of methylated sites. We found that global hypomethylation was present in the myoblast stage and is maintained during differentiation into the late myotube stage. Differential methylation analysis reveals an average hypomethylation across 2.5kb promoter regions of DUX4 target genes at both stages. Since mutant cells are derived from healthy control adult myoblasts unlike patient cell lines that are derived originally from patient differentiated muscle tissue in which *DUX4* and target genes might have already expressed, the hypomethylation at DUX4 promoters in proliferating FSHD mutant myoblasts suggests that low-level *DUX4* activity in undifferentiated myoblasts could be sufficient to allow activation of DUX4 target genes. Complementary long-read RNA-seq from the late myotube stage shows a correlation between hypomethylation at the *DUX4* promoter regions and an upregulation of DUX4 target genes. Although a majority of the promoter regions were hypomethylated, not all the transcripts at these promoters were significantly expressed, indicating that reduced methylation alone is insufficient for activation or that these targets may be poised but are not induced at this stage of muscle differentiation (sensitivity of DUX4 and some of the DUX4 target genes to differentiation has been reported (33)).

Long read enabled the identification of isoform-specific expression and methylation changes. Isoform-level expression analysis in FSHD is only beginning to be explored (72) and is particularly valuable for genes that are also implicated in other diseases. *DUX4* has been implicated in various cancers (73, 74) and one study has shown that in a few cancer cell lines, full-length *DUX4* mRNA expression originated from D4Z4 repeats in chromosome 4, both 4qA and 4qB alleles, and chromosome 10 that used alternative polyadenylation sites (73). The DUX4 target *SLC34A2* has been implicated in cancer (75, 76) and is a potential prognostic marker of oncological diseases (77). *SLC34A2* has 7 different isoforms and is currently being considered as a potential protein biomarker of FSHD (78, 79), however isoform level analysis for this gene has not been performed in the literature. Examining isoform-level expression may provide insights into its functional mechanisms and whether specific isoforms play unique roles in different disease contexts as seen in *DUX4* isoform expression in FSHD and cancer. Chromatin accessibility data for these transcripts would help validate their isoform specificity.

Other notable expressed isoforms were those associated with *DUXA* pseudogenes, particularly *DUXAP8* and *DUXAP9*. Strong evidence suggests DUXAP8 and DUXAP9 act as oncogenes promoting cancer malignancy (48–51). DUX4 reactivation in the context of cancers may promote cancer growth through activation of these lincRNAs. Interestingly, overexpression of *DUXAP8* and *DUXAP9* in renal cell carcinoma is linked to *COL1A1* and *COL1A2* upregulation (80). In FSHD, *COL1A1* and *COL1A2*, along with other collagen genes, are upregulated in patient muscle biopsies (21, 29). Therefore, these *DUXA* pseudogenes, which are more highly expressed than *DUXA* itself, may contribute significantly to the FSHD phenotype. Although COL1A1/2 are not significantly differentially expressed in our mutant myotubes *in vitro*, additional studies will be necessary to address their role using additional patient samples. Proper detection/mapping of these genes was only possible with long read sequencing. Expression of these genes are much more significant than their homolog DUXA in our FSHD mutant cells. This increased expression is accompanied by a greater reduction in promoter methylation compared with DUXA suggesting the need for future studies of the roles of these 2 pseudogenes in FSHD pathogenesis. Critically, characterization of these pseudogenes, as well as potentially other DUX4 target genes, must be carefully considered given the significant mapping discrepancies between Hg38 and Chm13. The CHM13 T2T human genome annotates the last missing 8% of the human reference genome, which contains highly repetitive and homologous regions, including macrosatellite repeats like D4Z4. Moreover, CHM13 resolves the gap in 4q35 (81), making it particularly valuable for studying FSHD. Since its release, studies investigating the disease locus utilized the complete human genome (69–71). Our analysis comparing the standard Hg38 reference with CHM13 using long reads suggests that the choice of reference assembly may be important not only for the DUX4 locus itself but also more broadly for accurately analyzing DUX4 target genes. Selecting the appropriate reference assembly is particularly important in disease studies, where accurate quantification of transcripts can influence our understanding of pathogenic mechanisms and the identification of potential biomarkers or therapeutic targets.

We performed *de novo* haplotype phasing to identify any isoform-specific expression and assess whether methylation levels exhibit allele-specific differences at D4Z4 locus, DUX4 target promoter regions and genomic repeats. Haplotype phasing adds another layer of resolution to help elucidate potential differences of DNA methylation and RNA transcription at different alleles and in our FSHD model, the permissive 4qA allele and the non-permissive 4qB allele. The entire D4Z4 locus was captured on individual reads, allowing characterization and direct measurement of methylation levels on each allele. The contraction of the FSHD mutant cell line created a *DUX4* promoter at DBET and hypomethylation was observed in both 4qA and 4qB alleles. The expression of full-length *DUX4* transcripts with different isoforms captured, however, are all mapped to 4qA consistent with our prediction. Hypomethylation at the disease locus extends genome-wide across all repeats, especially other satellite repeats like D4Z4 repeat arrays. This suggests that mechanisms that influence methylation at the disease locus might extend to other areas of the genome with similar structure. For the DUX4 target genes, phasing revealed monoallelic expression of three genes and allelic methylation patterns that are otherwise hidden in bulk analysis. A subset of promoter regions exhibiting differences in methylation between FSHD mutant cells and control are driven by changes on a single allele rather than the combined changes across both alleles. The use of a 2.5kb window to assess DNA methylation in promoter regions was necessary given the lack of annotated promoters for DUX4 target genes within ENCODE cCREs. However, closer examination across the 2.5kb window at various DUX4 target genes reveals smaller, discrete regions of differential methylation at other annotated regulatory elements or at potentially unannotated sites. This highlights the need for higher-resolution methylation profiling at the promoter itself rather than at broad regions upstream genes to more accurately assess allelic and isoform-specific methylation differences, which may require improved cCRE annotations.

What makes our allele-specific analysis particularly striking is that the comparison is performed against an isogenic control, indicating that the observed methylation changes appear to be the direct consequence of FSHD mutations. Although *de novo* phasing does not allow the definitive assignments of parental haplotypes to specific alleles and therefore cannot determine whether the more hypomethylated promoters comes from the same parent as the permissive allele, we observe differences when comparing allele-to-allele methylation (between H1 and H2) in the controls with the FSHD mutant cells This suggests that although the two alleles already differ in the control line, changes at the *DUX4* locus and/or SMCHD1 mutations caused additional allele-specific methylation changes in the FSHD mutant cells. This is further supported by principal component analysis of allele-specific methylation differences at genome-wide repeats where the biggest source of variation is driven by the alleles then the FSHD mutation. Given the genome-wide allelic epigenetic differences observed, it would be informative to characterize single-contraction edited cells lacking the SMCHD1 mutation to assess its extent of which SMCHD1 loss exacerbates hypomethylation. It should be noted, however, that SMCHD1 mutations introduced in adult myoblast line have no significant effect on either establishment or maintenance of DNA methylation status at D4Z4 despite the synergistic effect together with D4Z4 contraction on DUX4 target gene activation (33).

Overall our work leverages the advantages of Nanopore sequencing to further elucidate FSHD mechanisms in our FSHD cell line. This study builds upon our previous work (Kong et al., 2024) (33) that demonstrated that a subset of DUX4 targets functions within a feedback loop to reinforce and sustain their own expression. The promoter regions of the DUX4 target genes are already hypomethylated in the proliferating myoblast stage therefore may be contributing to the effective upregulation of these genes in differentiated myotubes. Highly expressed DUX4 target pseudogenes might also be missing noncoding disease modifiers in the feedback loop. Further studies are required to elucidate the biological functions of genes as candidate targets and to assess whether they should continue to be classified as pseudogenes or lncRNAs. Our work integrates long-read genomic and RNA sequencing to further define the epigenetic and transcriptomic landscape of FSHD.

## Limitations

Absence of parental information for the control cell line prevents the conclusion that the observed allelic hypomethylation occurs from the same parent that has the 4qa locus. Additional experiments using a trio is required along with chromatin accessibility information for robust characterization of the impact of the pathogenic allele on expression of DUX4 target genes and repeats. Lastly, immortalization of adult myoblasts may not accurately reflect gene regulation during embryonic genome activation.

## Data availability

Data is being deposited in dbGaP study accession phs002554. Direct-RNA isoforms dataset for swanViewer is available on Zenodo at DOI 10.5281/zenodo.18022278.

## Author contributions

A.M. and K.Y. designed the project. X.K. created and grew the cell lines as well as extracted the DNA and RNA. N.M. and E.T. built direct RNA libraries. N.M. also performed data analysis and edited the manuscript. A.M. and E.A. wrote swanViewer package. J.S.S built genomic libraries, performed data analysis, generated figures, and wrote and edited the manuscript with significant input from A.M. and K.Y.

## ACKNOWLEDGEMENTS

This work was funded by National Institute of Arthritis and Musculoskeletal and Skin Diseases (NIAMS) R01AR071287.

## Methods

### Culturing control and edited cell lines

Immortalized skeletal myoblast control cells and double mutant (DEL_SM) cells were cultured in growth medium consisting of high-glucose DMEM (Gibco) supplemented with 20% FBS (Omega Scientific, Inc.), 1% Penicillin-Streptomycin (Gibco), and 2% Ultrasor G (Crescent Chemical Co.). The double mutant line, referred to as DEL9_SM_A in our previous paper, was generated from the control cells using the Alt-R CRISPR-Cas9 genome editing system (IDT) as described (Kong X. et al., iScience, 2024) (33). This line carries a homozygous deletion in the SMCHD1 gene and a contracted 4qA allele containing only a single D4Z4 repeat.

Upon reaching approximately 80% confluence, differentiation was induced by switching to high-glucose DMEM supplemented with 2% FBS and 1% ITS (Thermo Fisher Scientific). The differentiation medium was refreshed daily. Differentiating cells were collected on day 12 for control cells and day 14 for the double mutant cells, respectively.

### Direct whole-genome long-read sequencing

Multiple Nanopore libraries were built for myoblast and myotube replicates for both control and DEL9_SM for multiple loadings on the Nanopore run. Each library was built with 1.4 µg input of labeled DNA using the Ligation Sequencing Kit V14 (ONT, cat. #SQK-LSK114) and NEBNext Companion Module for Oxford Nanopore Technologies Ligation Sequencing (NEB, cat. #E7180). AMPure XP beads (Beckman Coulter, cat. #A63881) were used for the clean-up and the Long Fragment Buffer (LFB) from the Ligation Sequencing Kit was used during the wash steps. After the last elution, beads were incubated in Elution Buffer overnight at 4°C on a vertical rotating mixer at 2 rpm. The next day, beads were pelleted on a magnet, libraries were recovered, and concentration was measured using the Qubit dsDNA HS Assay Kit (Thermo, cat. #Q32854).

### Sequencing direct DNA libraries

Since multiple libraries were built per sample, each library was used as one load on a PromethION R10.4.1 Flow Cells flow cell (ONT, cat. #FLO-PRO114M) at 400-450 total ng per library. Flowcell priming mix with Bovine Serum Albumin (BSA) (0.2 mg/ml final concentration) and library mix were prepared according to the protocol. Libraries were sequenced on a PromethION 2 Solo with the Oxford Nanopore’s MinKNOW sequencing software (v24.02.16) at 400 bps. Libraries were loaded once each day after a flow cell wash (ONT, cat. #EXP-WSH004) until the end of flow cell life (5-7 days).

### Sequencing direct RNA libraries

Nanopore libraries were built from myotube replicates for both control and DEL9 SM for single loading on the Nanopore runs. Each library was built with an average of 700ng input of total RNA that was eluted in 10uL of nucleus-free water using Direct RNA sequencing kit (ONT, cat. #SQK-RNA004). Instead we used 2.5uL of 10 mM dNP solution (NEB, cat. #N0447) and 2.5 uL of 5x Induro RT reaction buffer (NEB, cat. #M0681) when making the reverse transcription master mix. Libraries were recovered, and the concentration was measured using the Qubit fluorometer DNA HS assay (Invitrogen, cat. #Q32856).

### Processing of Nanopore data

Raw Nanopore reads from both DNA and RNA were processed using Dogme (37) v1.2.2. This nextflow pipeline basecalled pod5 files using dorado v1.0.0. Reads were mapped using minimap2 v2.28 and samtools v1.15.1. The mapping preset used for genomic reads was -ax lr:hq and for RNA was -ax splice:hq (for the full list of parameters, see the package documentation). Modifications were extracted from bam files into bedMethyl files using modkit v0.5.0 and the transcripts were quantified using LR-Kallisto and bustools v0.51.1 and v0.43.2 respectively. For RNA reads, modification probability threshold was set at 0.9. For genomic reads, modkit sets a modification probability threshold by sampling 10,000 reads and calculating the modification probability threshold to filter out the 10% lowest confidence modification calls.

### Differentially methylated site calling

Dispersion Shrinkage for Sequencing data (DSS) (82) v2.50.1 was used for calling DMSs. DSS is based on a Bayesian hierarchical model to estimate and shrink CpG site-specific dispersions, then conduct Wald tests for detecting differential methylation. Input for DSS was modified from the preprocessed methylation bed file. Bedmethyl file was filtered for a minimum coverage set at 3 reads and the maximum coverage was set individually for each sample at the 99th percentile coverage. Modifications were subset as their own input files. These filtered bed files were further edited to only include the required fields by DSS: chromosome number, genomic coordinate, total number of reads, and number of reads showing methylation. To optimize resource usage and time, bed files were further split by chromosome. The DMLtest function is used to estimate mean methylation levels for all CpG sites, estimate dispersions at each CpG site, and conduct Wald tests. The output is an R object, which is passed to the callDMR function where regions with many statistically significant CpG sites are identified as DMSs. We applied a 0.05 p-value threshold. The output tsv files for each chromosome were concatenated together for each modification.

### Measuring methylation level of regions

Promoter regions were calculated from a gencode annotation file using a custom script. Transcription start site was determined using strand orientation and start position. Then two thousand base pairs upstream and five hundred base pairs downstream was calculated, ensuring that no region exceeds chromosome boundaries by factoring in chromosome length. Transcript isoforms that overlap less than 50% from each other are kept as separate promoter regions. Methylation level differences exceeding two standard deviations at the promoters of DUX4 target genes in edited cells compared with controls were considered significant. RepeatMasker Repetitive Elements track and Mappability track with Single-read and multi-read mappability track was downloaded from UCSC Genome Browser and converted from a BigBed to a Bed file using the bigBedToBed utility from the UCSC Genome Browser. To score and calculate percent methylation of a region, the bed file was passed through the modkit dmr pair subcommand (v0.4.3) with a pair of sample bedmethyl files. Promoter regions exhibiting no change between FSHD and controls were regions with a mean methylation difference change 5% or less. The Integrative Genomics Viewer (IGV) (83) was used to visualize methylation.

### Phasing genomic reads

All the edited samples and their isogenic controls were merged into one bam file with round 100x coverage for generating a VCF file using Clair3 (84). Clair3 v1.1.1 was used with the following parameters: --platform=“ont” --model_path=r1041_e82_400bps_sup_v500 --use_whatshap_for_final_output_phasing --enable_long_indel. Phasing was done using WhatsHap (85) v2.7 using default parameters. Promoter regions with fewer than three reads of coverage due to incomplete phasing were excluded, resulting in the removal of eighteen transcripts from fifteen FSHD genes, including RFPL2-203, which was a significantly hypomethylated gene before phasing.

### Reconciling novel isoforms across different samples

After running the Dogme pipeline for RNA annotation, each sample has its own set of novel isoforms. We developed reconcileBams.py to generate a unified consensus transcriptome by aggregating novel read support across all samples. This process reconciled novel isoforms by merging incomplete splice matches (ISMs) into representative transcripts and correcting strand assignment errors. To ensure robustness, novel isoforms were rigorously filtered (retained only if >0.1 TPM in 2 samples) to produce a final high confidence GTF and consistently tagged BAM files as well as an abundance file of known and novel isoforms for downstream analysis. The reconcileBam.py script is available within the Dogme repository at: https://github.com/mortazavilab/dogme/tree/main/scripts.

### Downstream analysis of RNA expression level and isoform usage using swanViewer

We developed the swanViewer web application using the Streamlit framework to facilitate interactive exploration of transcript isoforms (86) and differential expression (Fig. S3). The reconciled transcriptome GTF and abundance matrices were first pre-processed using the helper script processSwan.py to optimize performance and reduce initialization time. This utility constructs a SwanGraph data structure, integrating reference annotations with the consolidated novel isoforms and serializes it, along with experimental metadata, into binary Python pickle (.p) files. These pre-computed pickle files allow the viewer to rapidly load the dataset, bypassing the computationally intensive graph construction step during interactive sessions. The app is designed to interactively analyze gene and transcript level differential expression and isoform analysis as well as visualize transcript usage for individual genes. The swanViewer application including a docker image is available at: https://github.com/mortazavilab/swanViewer.

## Supplementary Tables

- **Table S1: DEGs Table of DUX4 target genes mapped to Chm13**.
- **Table S2: DEGs Table of DUX4 target genes mapped to Hg38**.

## Supplementary Figures

- **Fig. S1: Methylation is not reduced at unedited isogenic control**.
- **Fig. S2: Subset of methylation difference at DUX4 target gene with isoform-specific expression**.
- **Fig. S2: Screenshot of swanViewer interface**.

