## Supplemental Figures for "Genome-wide analysis of FSHD cell lines using Nanopore sequencing reveals allele-specific differences at DUX4 target genes and complex repeats"

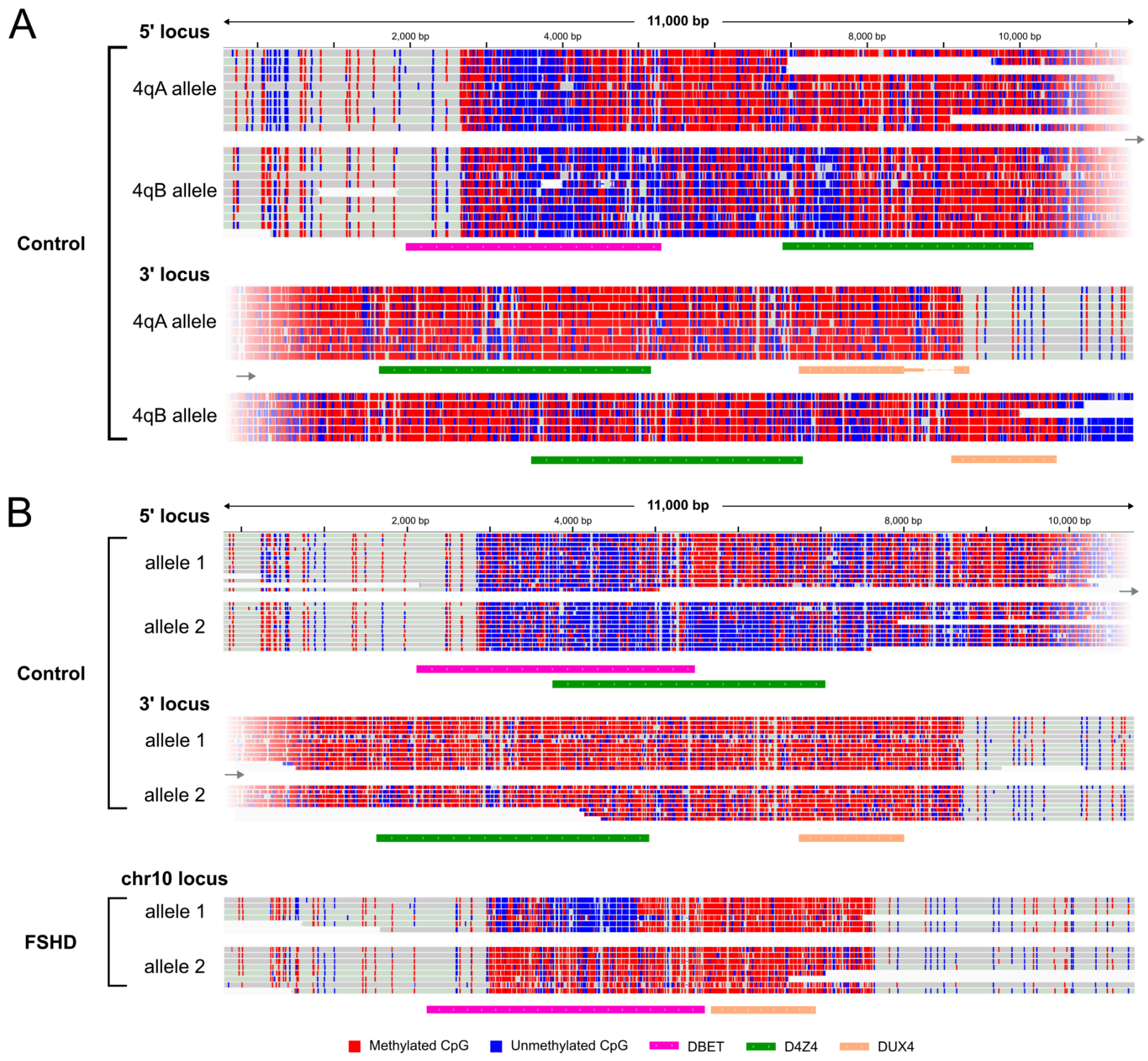


**Fig.S1. Methylation is not reduced at unedited isogenic control. A.** Genomic reads of the isogenic control at the D4Z4 locus on 4q35. The top two tracks show the 5' end of the D4Z4 locus with DBET and the first D4Z4 repeat unit. The last two tracks show the 3' end of the D4Z4 locus with the last two D4Z4 repeat units. **B.** Genomic reads of the isogenic control and FSHD mutant at the D4Z4 repeats on 10q. The top four tracks show reads from isogenic control mapped to 10q (split by 5' and 3' end) and two bottom tracks show the contracted 10q in FSHD mutant cells.

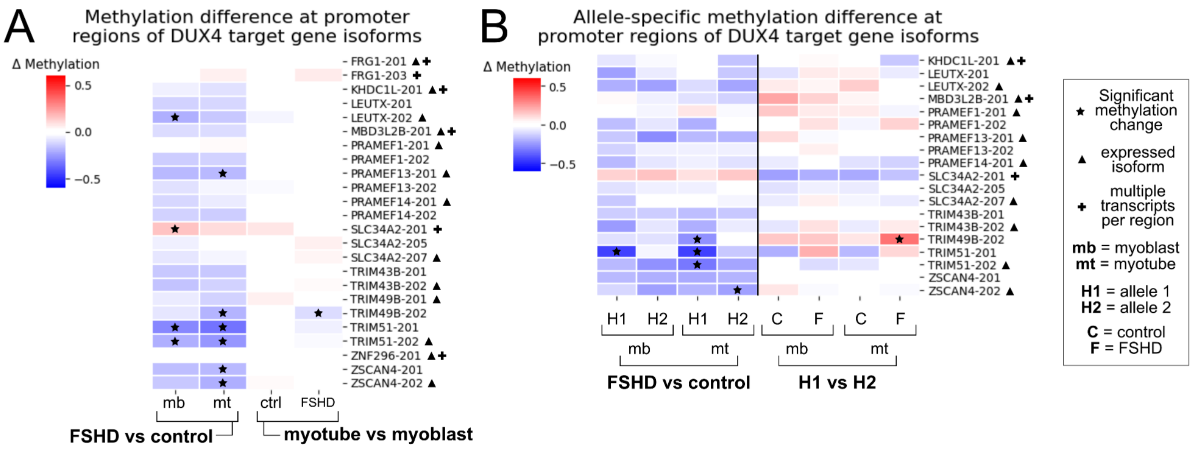


**Fig. S2. Subset of methylation difference at DUX4 target gene with isoform-specific expression. A.** Methylation difference at DUX4 target promoter regions between (left to right) FSHD mutant and control myoblasts, FSHD mutant and control myotubes, control myotubes and myoblasts, and FSHD mutant myotubes and myoblasts. Significant methylation, isoform expression and promoter regions with multiple isoforms are annotated with a star, triangle, or a plus (respectively). **B.** Allele-specific methylation difference (H1 vs H2) at DUX4 target gene promoter regions between (left to right) FSHD mutant H1 and control H1 as well as FSHD H2 and control H2 at myoblast stage (mb) and myotube stage (mt). H2 and H1 comparison in control (C) and FSHD (F) myoblast (mb) and myotubes (mt) stages. Significant methylation, isoform expression and promoter regions with multiple isoforms are annotated with a star, triangle, or a plus (respectively).

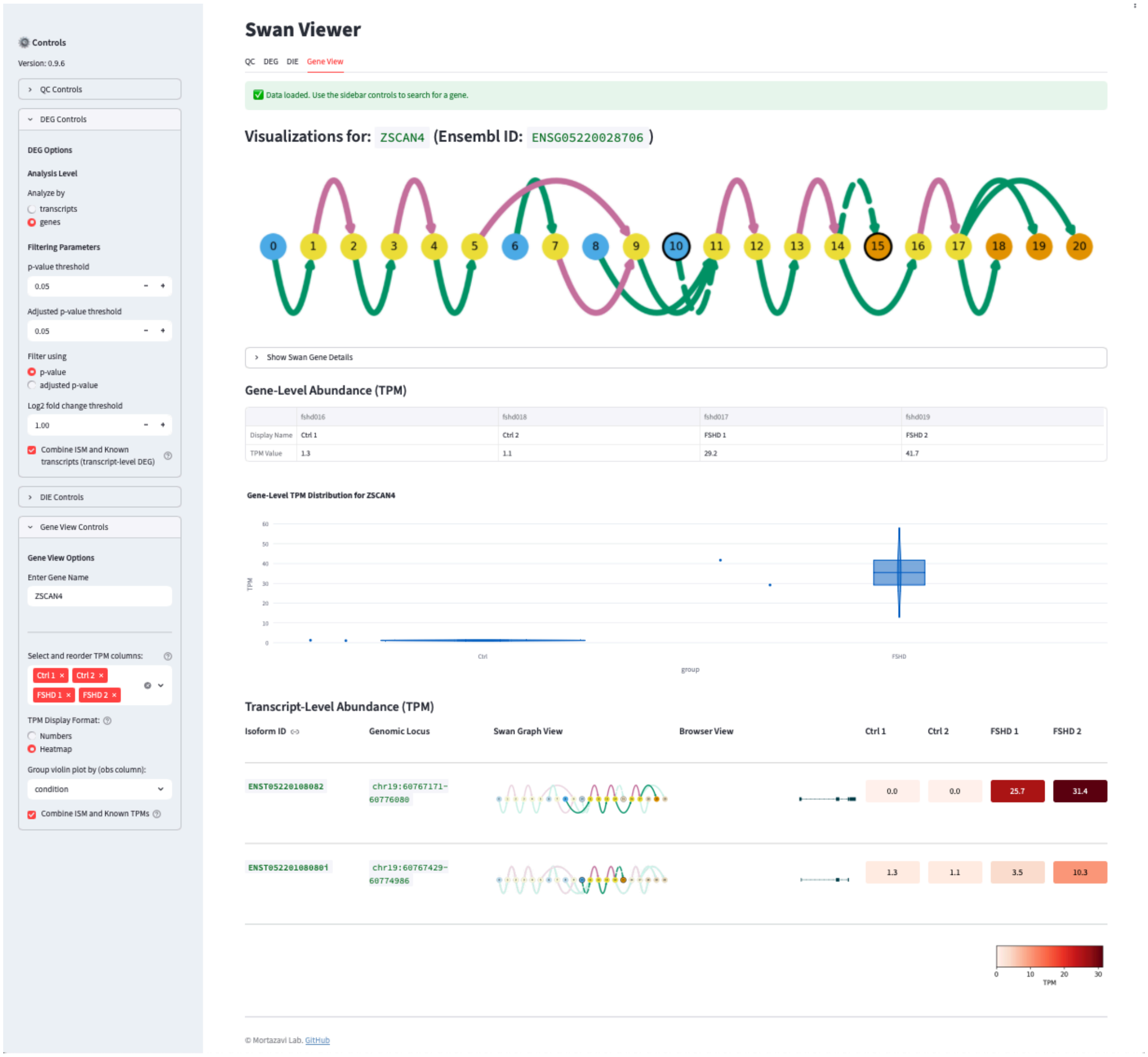


**Fig. S3. Screenshot of swanViewer interface.** Shows *ZSCAN4* transcript TPM and models in the "Gene View" tab.
